# Dynamics of Sox2 expression during rat germ cell development and its relationship with emergence of spermatogonia

**DOI:** 10.1101/558015

**Authors:** Ticiana Volpato de Oliveira, Renato Borges Tesser, Marina Nunes, Taiza Stumpp

## Abstract

The gonocytes represent a specific phase of male gem cell development that precedes spermatogonial stem cell differentiation. Here, we describe the expression of Sox2, an OCT4 partner, during rat germ cell development. Our hypothesis is that SOX2 has a cytoplasmic role during gonocyte-to-spermatogonia transition. Male rat embryos and testes were submitted to the analysis of Sox2 expression. *Sox2* was detected in germ cells from 14 days post-conception (dpc) to 8dpp. SOX2 was present in 14dpc and 15dpc embryos and absent at 17 and 19dpc; however, it did not show direct correlation with mRNA. SOX2 labelling was detected after birth and its expression increased from 1dpp to 5dpp. SOX2 was localized in the cytoplasm and showed a granulated pattern similar to P-bodies. Indeed, GW182/SOX2 and LIN28/SOX2 double-labelling showed that SOX2 partially co-localized with the P-bodies components GW182 and LIN28. At 8dpp SOX2 was detected in the nucleus and/or in the cytoplasm of spermatogonia, whereas at 25dpp it was detected in the nucleus of rare spermatogonia. This suggests that SOX2 localization changes during gonocytes to spermatogonia transition.

## 2. INTRODUCTION

Spermatogenesis is a very dynamic and controlled process that culminates with the differentiation of the sperm. The maintenance of spermatogenesis depends on the proliferation and differentiation of the spermatogonial stem cells (SSC) that reside in specific compartments of the seminiferous epithelium called niches. Although the presence of stem spermatogonia in the testis has been reported more than 40 years ago (1), very few is known about their biology and about the mechanisms that drive their differentiation from the gonocytes. In addition, it is not even clear what is the identity of these cells.

The SSC can be considered a special type of stem cell since they express a variety of embryonic stem cell markers *in situ* such as Tex19.1. (2), Sall4 (3), Lin28 (4, 5) and Oct4 (6). This places the SSC as an intermediary between embryonic and adult stem cells (7, 8, 9) and makes these cells very interesting and promising, since they can be reprogrammed to a pluripotent state *in vitro* without any genetic manipulation.

The SSC differentiate from the gonocytes, which represent a phase of male germ cell development that comprises the moment when they reach the gonads and the first week of postnatal phase (10). During this phase they go from a mitotic phase to a quiescent (or mitotic arrest) phase and back to a proliferative state, when they migrate to the basis of the seminiferous cords and differentiate into spermatogonia (10). The quiescence period is characterized by the downregulation of proliferation markers and of pluripotent markers present in primordial germ cells (PGC) such as Oct4 in rats (11, 12, 13) and *Sox2* and *Nanog* in mice (12). In humans Oct4 and Nanog were detected in primordial germ cells (14), but the downregulation of these genes does not seem to lead to a quiescent condition (13). Contrasting with mice, Sox2 has not been detected in human primordial germ cells (PGC) (15, 16, 17).

In rats, by the end of quiescence and at the moment of proliferation resumption, OCT4 protein is upregulated and become a marker of undifferentiated spermatogonia from this stage on (11). In mice Oct4 and Sox2 has been detected at mRNA levels in perinatal and adult germ cells (18, 19), but only OCT4 was detected at protein level (18). On the other hand, Nanog has not been detected after birth (20). In humans, the data about Oct4 and Nanog expression is controversial: some studies detect Nanog and Oct4 in the adult testis (21) whereas others do not detect these markers at any time after birth (14). Conversely, Sox2 expression has not been detected in pre-pubertal and adult testes (15, 16, 17). However, to our knowledge, the expression pattern of Sox2 and Nanog during rat germ cell development has not been explored.

We have previously suggested that germ cell development in rats is different from mice and seems to be more similar to humans (22). Thus, we decided to investigate Sox2 expression in rat germ cells at PGC stage, during mitotic arrest and at proliferation resumption, which coincides with the transition from gonocytes to spermatogonia. Our data suggest that Sox2 expression varies according to the phase of germ cell development and shows different dynamics, especially at the protein level, from that described for mice and humans. We also suggest that Sox2 seems to be an important factor for SSC emergence.

## 3. MATERIALS METHODS

### 3.1 Materials

All primers for conventional and quantitative PCR were obtained from IDT Technologies (Sintese Biotecnologia, Brazil). The antibodies were obtained as follows: anti-SOX2, from Abcam (Cat. ab59776), anti-LIN28 from Novus Biologicals (Cat. IMG6550A), anti-GW182 from Santa Cruz Biotechnology (Cat. sc66915) and anti-*ß*Actin from Cell Signalling (Cat 4970S). The kits for immunohistochemistry (DAKO Detection System) and Western blot (Western Dot® 625) detection were obtained from DAKO (Cat. K0690) and ThermoFisher Scientific (Cat. W10142), respectively. The secondary antibodies for immunofluorescence were obtained from Millipore (Cat. 12-507), Vector Laboratories (Texas Red Cat. TI-1000) and Abcam (Alexa Fluor 5894® ab150116). The protease inhibitor cOmplete™, EDTA-free Protease Inhibitor Cocktail (Cat 11873580001) and all reagents used for the buffers were obtained from Sigma Aldrich/Merck or ThermoFisher Scientific.

### 3.2 Animals and Tissue Preparation

Pregnant Wistar rats (*Rattus norvegicus albinus*) were obtained from the Laboratory of Developmental Biology (UNIFESP, Sao Paulo, Brazil). The animals were kept in plastic cages under a 12–12 hours light/dark cycle at 23-25°C. Food and water were allowed *ad libitum*. Pregnancy was detected by the presence of sperm in the vaginal smears (1dpc).

### 3.3 Embryo and testis collection

The dams were euthanized by analgesic/anaesthetic (xylazin/ketamin, 10 mg/Kg and 100 mg/Kg, respectively) method. The embryos were collected at 14, 15, 17 and 19dpc and fixed in Bouin’s solution for immunohistochemistry. Embryo sexing was performed by visual inspection of the gonads. For testis collection, 1dpp, 5dpp, 8dpp and 25dpp rats were submitted to euthanasia through decapitation (1dpp, 5dpp and 8dpp) or by analgesic/anaesthetic (xylazin/ketamin, 10 mg/Kg and 100 mg/Kg, respectively) method (25dpp). The embryos from each age were obtained from five different mothers and the testes from 5 individuals to guarantee sample variability. The experiments were carried out according to the rules of the local committee for animal care (CEP No. 0251/12) and the National Institutes of Health (NIH) guide for the care and use of Laboratory animals.

### 3.4 Immunolabelling

Embryos (15 and 17dpc, n=6) or testes (19dpc, 1dpp, 5dpp, 8dpp and 25dpp, n=5) were fixed in Bouin’s liquid and processed for paraffin embedding. Four cross-sections (6 μm-thick) were obtained from each embryo or testes and submitted to the labelling of SOX2, LIN28 and GW182. The sections were dewaxed in xylene, hydrated and submitted to heat antigen retrieval using citrate buffer (pH 6.0.) for 10 minutes. The slides were treated with 5% BSA and incubated with the primary antibodies anti-SOX2 (1:250, Abcam - ab59776), anti-LIN28 (1:250, Novus Biologicals/Imgenex - IMG6550A) and/or GW182 (1:200, Santa Cruz - sc66915) overnight at 4°C. For immunohistochemistry, the slides were washed in PBS (0.0.5 M, pH 7.2.) 3x and incubated with the secondary antibody (DAKO Detection System - K0690, USA). The slides were washed again in PBS and then incubated with the streptavidin-peroxidase (DAKO Detection System - K0690, USA). The reaction was revealed with DAB (K3468, DAKO, USA) and the nuclei were stained with Harris Hematoxilyn. For SOX2/GW182, SOX2/LIN28 and LIN28/GW182 double-labelling, FITC, Alexa and Texas Red-conjugated secondary antibodies were used. DAPI was used for nuclear staining. Negative controls (primary antibody omission) were performed for all reactions.

### 3.5 RT-PCR

The total RNA from embryo gonads (n=3) at 15, 17 and 19dpc and from 5dpp and 8dpp testes (n=3) was isolated using TRIzol reagent (Invitrogen®). The cDNA was obtained using the Superscript IV Reverse Transcriptase (Invitrogen®). Quantitative RT-PCR (RT-qPCR) was performed using PowerUp SYBR Green Master Mix (Applied Biosystems) and specific primers for *Sox2* (Table 1). Conventional RT-PCR was performed to check the presence of *Oct4* and *Nanog*, which are the other two pluripotency factors of the “Pluripotency Triplet”. *βActin* was used as reference gene for the conventional and quantitative PCR. The primer sequences are shown in Table 1.

**Table 1.**
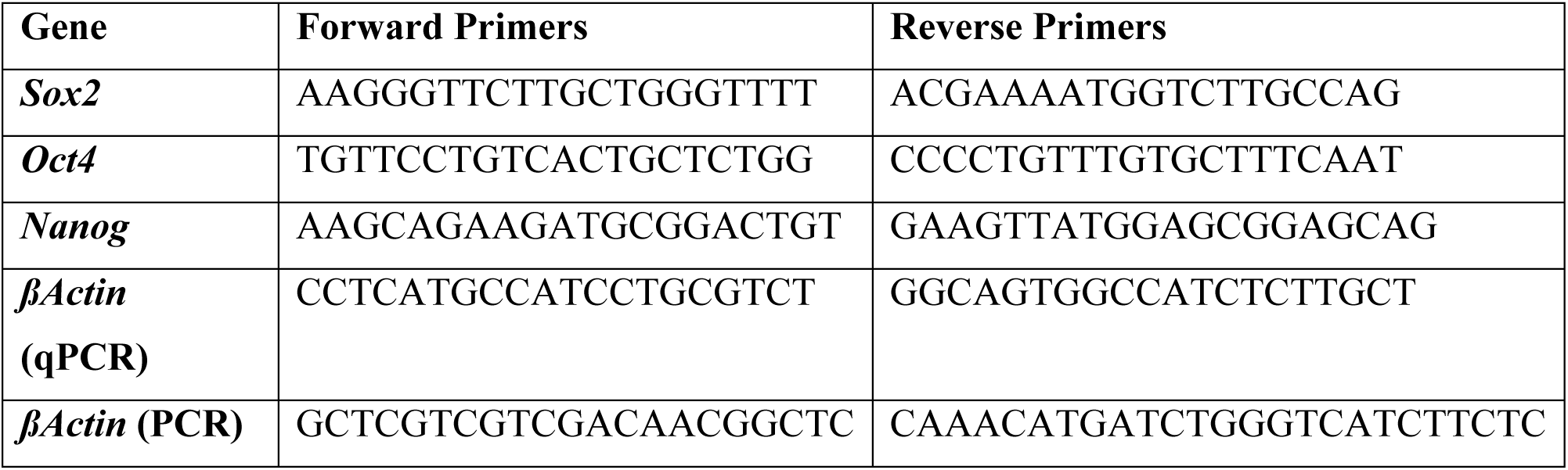
Sequences of primers used for RT-qPCR (*Sox2* and *βActin*) and RT-PCR (*Oct4, Nanog* and *Gapdh*).

### 3.6 Western blot

To confirm SOX2 expression in the quiescent gonocytes, the testes were collected at 1dpp and submitted to lysis buffer (50mM Tris-HCl,25mM NaCl, 1% Triton X-100) and sonication. The protease inhibitor cOmplete™, EDTA-free Protease Inhibitor Cocktail (Merk) was added immediately. The protein extract was separated by SS/PAGE in an 8% polyacrylamide gel and the bands were transferred to a nitrocellulose membrane. The membrane was incubated overnight with the same anti-Sox2 antibody used for the immunohistochemistry reactions (1:200, Abcam - ab59776). The secondary antibody incubation and bands revelation was performed using the Western Dot® 625 (Cat. W10142, Invitrogen, Thermo Scientific™) in an E-Gel™ Imager System with Blue Light Base (ThermoFisher Scientific). *ß*-Actin was used as reference.

## 4. RESULTS

### 4.1. RT-qPCR

To investigate *Sox2* expression during gonocytes quiescence, we performed RT-qPCR. *Sox2* expression was detected at all ages, but the level of expression decreased from 15dpc to 8dpp (Figure 1A). This data do not exactly match the immunohistochemistry analysis in which SOX2 was detected at 15dpc and then from 1dpp to 8dpp (Figure 1A). This inconsistency could be explained by the limited detection capacity of the techniques used or may simply reflect a biological process in which the expression of Sox2 at mRNA or protein levels SOX2 is regulated according to the germ cell development.

**Figure 1.**
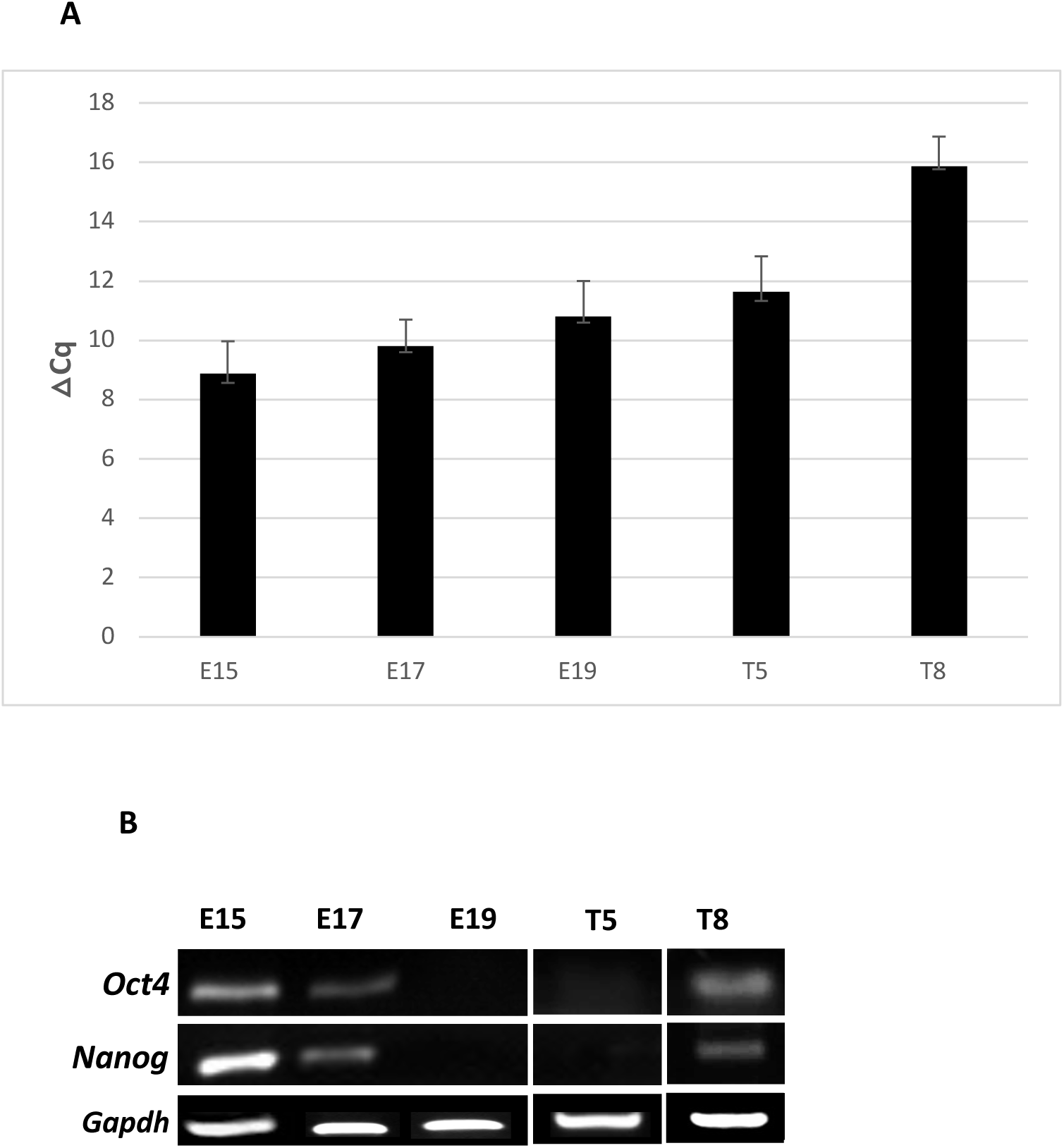
Sox2 expression in 15dpc and 19dpc embryo gonads and 5dpp and 8dpp testis. A: Sox2 was detected at all ages but showed a decrease in the expression level at 8dpp (**p<0.001). B: The conventional PCR shows that *Nanog* and *Oct4* are detected at 15dpc, 17dpc and 8dpp, but not at 19d0c and 5dpp.

To check for the presence of the other two members of the pluripotency triplet, the expression of *Oct4* and *Nanog* was investigated at the same ages by conventional PCR. The expression of these genes was detected at 15dpc, 17dpc and 8dpp, i.e., during gonocyte or spermatogonia proliferation, but not at 19dpc and 5dpp, when gonocytes are quiescent (Figure 1B).

### 4.2 Immunohistochemistry

During rat male germ cell development SOX2 expression was clearly dependent on the stage of germ cell development. SOX2 expression was detected in the gonocyte nucleus at 14dpc (Figure 2A) and 15dpc (Figure 2B). SOX2 was not detected at 17dpc (Figure 2C) and at 19dpc (Figure 2D), when germ cells are quiescent (23, 11). This suggests that SOX2 is downregulated as male germ cells enter mitotic arrest. SOX2 labelling was detected again at 1dpp and was localised in the gonocyte cytoplasm (Figure 3A). At 5dpp the SOX2 labelling in the cytoplasm of the gonocytes was more abundant that at 1dpp (Figure 3B). At both ages SOX2 labelling showed a granulated pattern very similar to the P-bodies, which are subcellular compartments responsible for RNA-processing. At 8dpp (Figure 3C to 3E) SOX2 labelling was detected in pre-spermatogonia nuclei (Figure 3C) or cytoplasm (Figure 3D), suggesting that this protein starts to be transported to the nucleus at this age, coinciding with OCT4 upregulation (11). At 25dpp SOX2 was detected exclusively in the nucleus of very rare spermatogonia (Figure 3E), suggesting that SOX2 could be a good indicator of the SSC in the rat testis.

**Figure 2.**
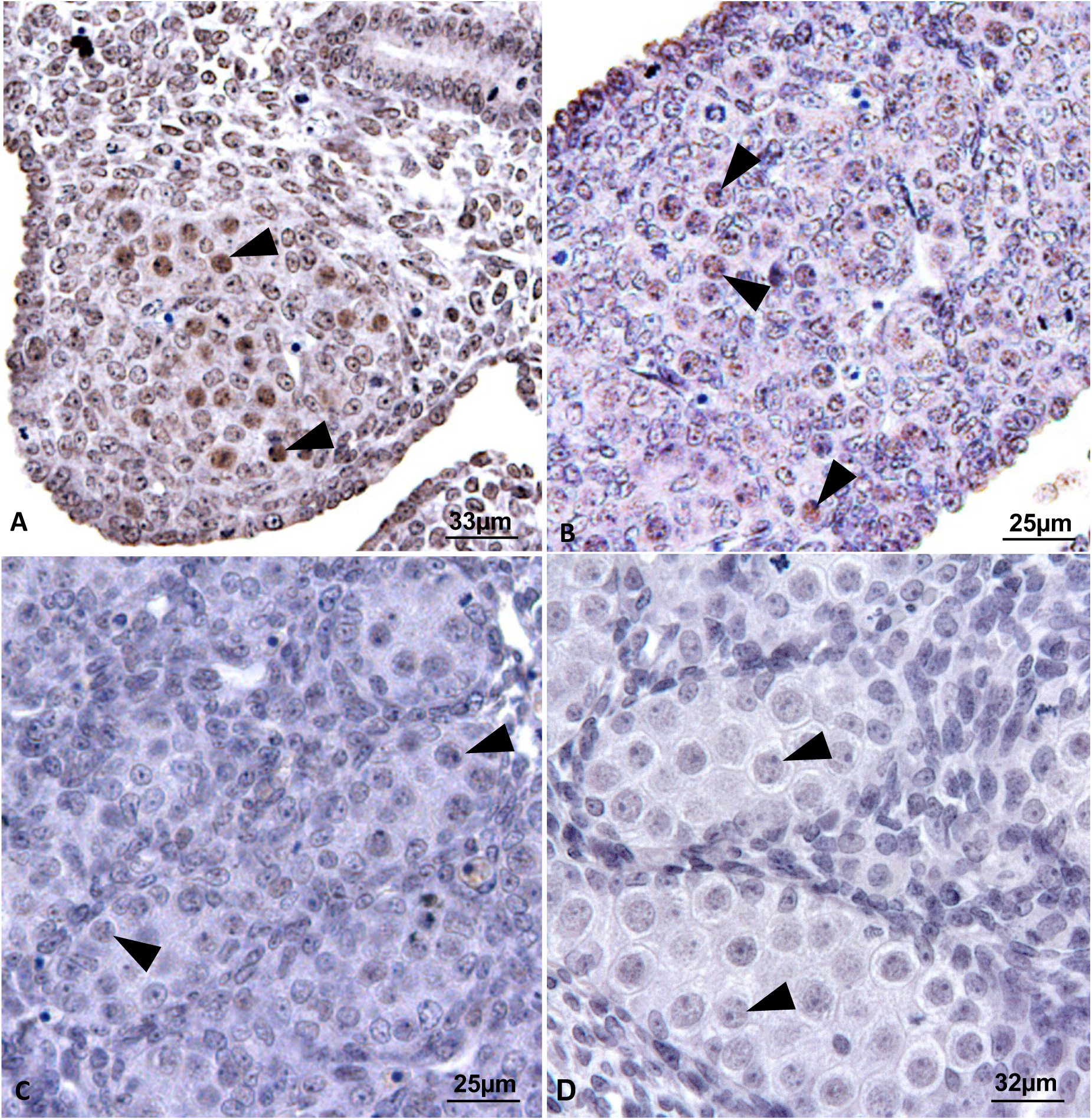
SOX2 immunolabelling in primordial germ cells of rat embryos at 14dpc, 15dpc, 17dpc and 19dpc. SOX2 was detected in PGC (arrowheads) at 14dpc (A) and 15dpc (B) but not at 17dpc (C) and 19dpc (Figure D).

**Figure 3.**
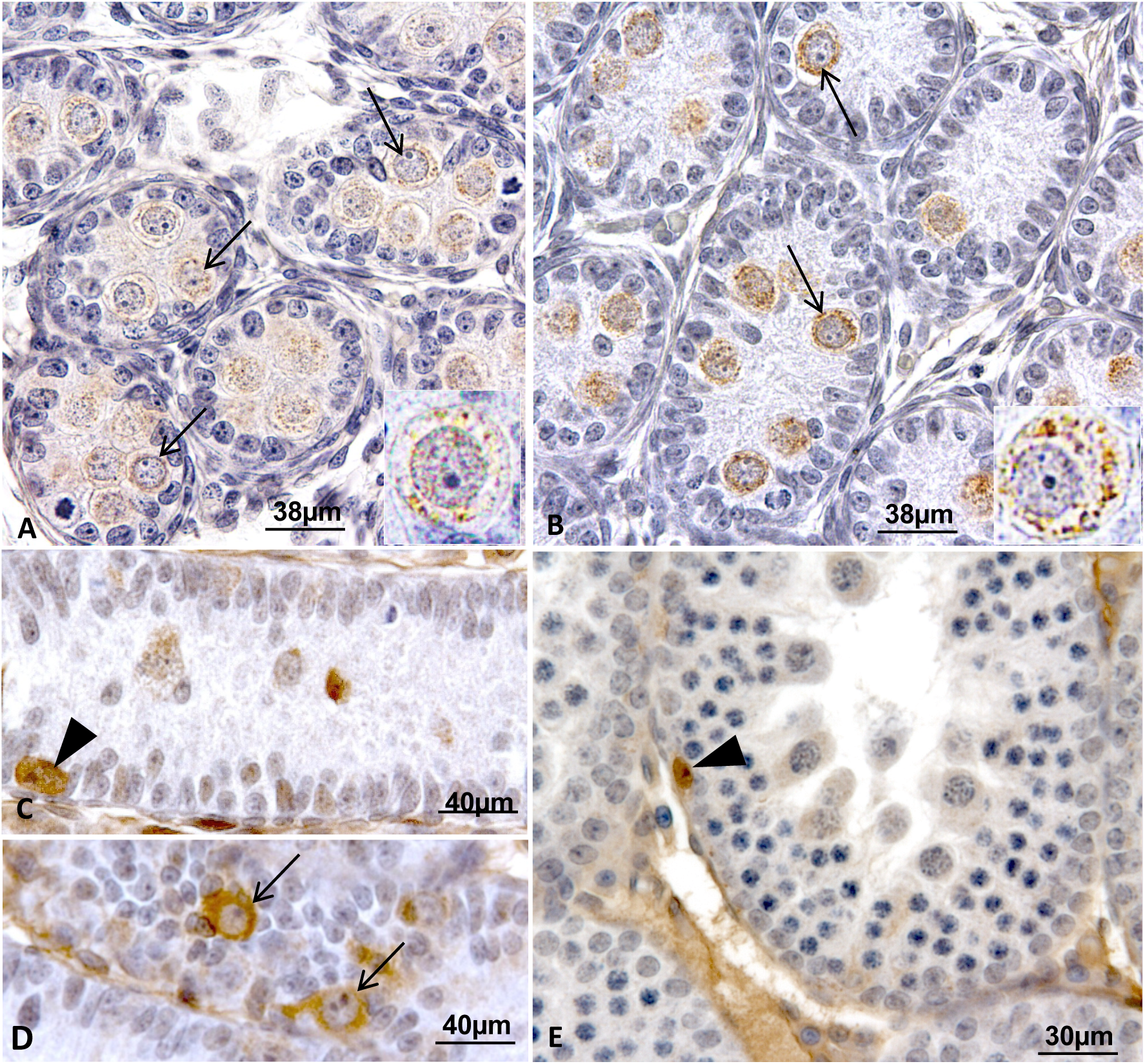
SOX2 immunolabelling in post-natal (1dpp, 5dpp and 8dpp) and pre-pubertal (25dpp) testes. SOX2 was detected in gonocyte cytoplasm (arrows) at 1dpp (A) and 5dpp (B). SOX2 labelling shows a granulated pattern and is less abundant at 1dpp than at 5dpp. The insets show the detail of SOX2 subcellular localization in gonocyte cytoplasm. At 8dpp (C and D) SOX2 was detected in the nucleus (arrowhead) or in the cytoplasm (arrows) of pre-spermatogonia. In 25dpp (E) testis SOX2 was present only in the nucleus of rare undifferentiated spermatogonia (arrowhead).

To infer about a possible function for SOX2 in RNA processing in the gonocytes, 5dpp testes were submitted to GW182 labelling, which is considered a marker of the P-bodies. The age of 5dpp was chosen because SOX2 labelling was more abundant that at 1dpp. GW182 labelling (Figure 4A) was very similar to SOX2 labelling (Figure 4B), with granulated pattern throughout the cytoplasm. On the other hand, contrasting with SOX2, GW182 was not specific to gonocytes and was also detected in Sertoli cell cytoplasm (Figure 4A). LIN28, a spermatogonial stem cell marker (4, 5), has been shown to be a component of the P-bodies (Balzer and Moss 2007) and to interact with SOX2 in embryonic stem cells (24). Thus, we investigated LIN28 labelling in post-natal gonocytes. Indeed, LIN28 labelling pattern was very similar to that observed for SOX2 and GW182 (Figure 4C).

**Figure 4:**
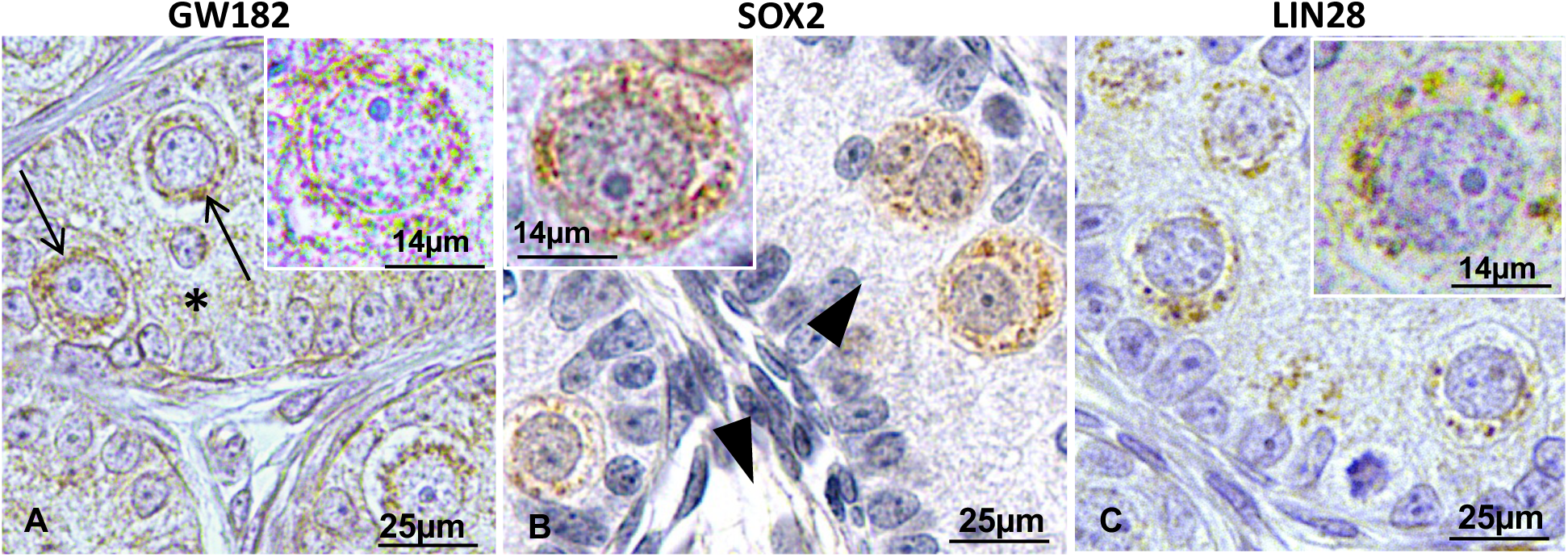
Comparison of GW182 (A), SOX2 (B) and LIN28 (C) immunolabelling in 5dpp testis. The three proteins show a similar granulated pattern in the gonocytes

To confirm whether SOX2 was localized in the P-bodies, we performed the double-labelling of SOX2 and GW182 as well as SOX2 and LIN28 in the 5dpp testes. We found that SOX2 and GW182 labelling partially coincides in 5dpp quiescent gonocytes (Figure 5), suggesting that some of SOX2 protein localizes to P-bodies in these cells. We also observed a partial co-localization of SOX2/LIN28 in the gonocyte cytoplasm (Figure 5), reinforcing that SOX2 might be part of P-bodie, since LIN28/GW182 double-labelling confirmed LIN28 localization to P-bodies (Figure 5

**Figure 5.**
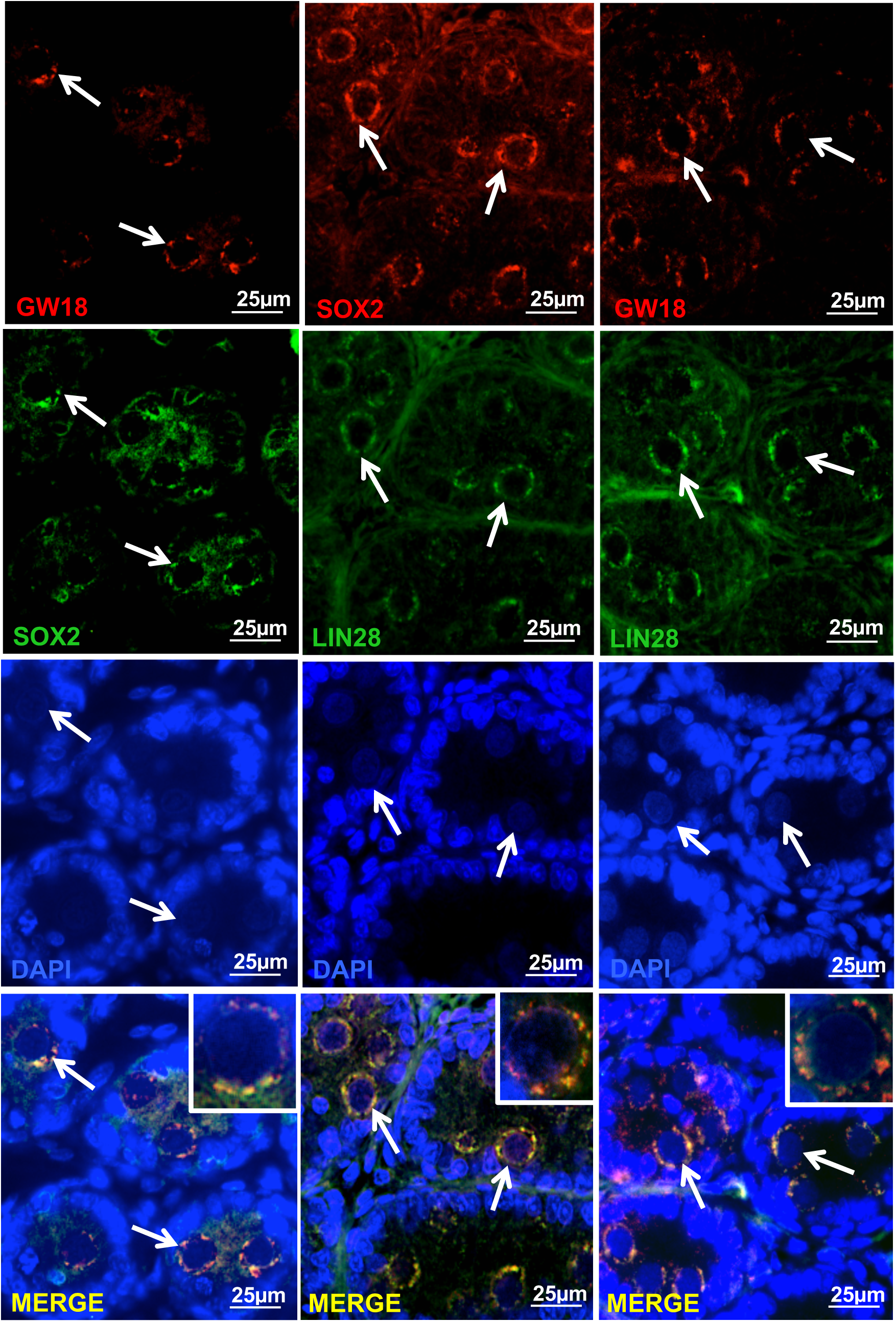
SOX2/GW182, SOX2/LIN28 and LIN28/GW182 double-labelling. SOX2 partially co-localizes with GW182 and Lin28, whereas LIN28 and GW182 show greater co-localization.

### 4.3 Western Blot

To confirm the expression of SOX2 during the quiescence phase, we performed a Western blot. Two specific bands were detected in the testicular extract of 19dpc testis: one of around 100KDa and another of around 60KDa (Figure 6). The expected band for SOX2 is of around 38 to 43KDa. However, considering that the reaction was very specific and that the antibody is well validated, we suggest that the two bands might correspond to the detection of complexes of SOX2 and other proteins that might interact with it.

**Figure 6.**
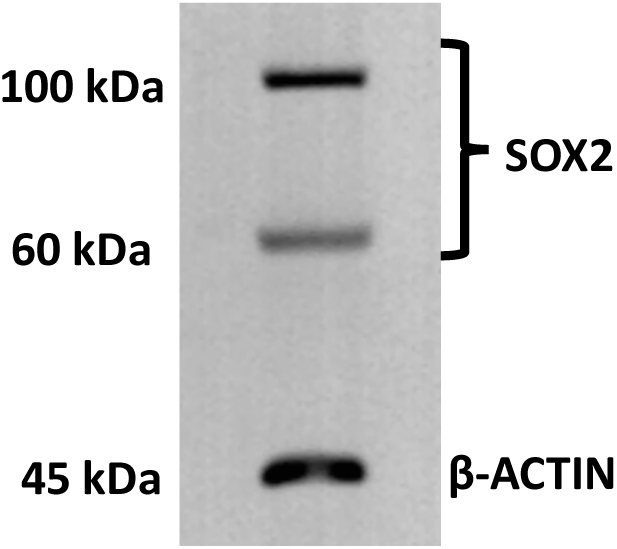
SOX2 expression in post-natal rat testicular homogenates. Representative immunoblot demonstrating the presence of SOX2 at 1dpp, confirming SOX2 presence in quiescent gonocytes.

## 5. DISCUSSION

Spermatogonial stem cells (SSC) are the male germline stem cells and are responsible to maintain spermatogenesis throughout life. Spermatogonia emergence is preceded by a quiescent period in which germ cells do not proliferate or die, but become distributed along and across the seminiferous cords (11). Male germ cell quiescence in mice and rats seems to involve the downregulation of the pluripotent marker OCT4 (22, 11) and of proliferation markers (25). Here we observed that Sox2 seems to follow a similar pattern of that observed for OCT4 expression as germ cells go towards quiescence during embryonic phase. Similarly, Oct4 and Sox2 downregulation at both mRNA and protein levels as PGC enter mitotic arrest has been also reported for mice (12). On the other hand, these data contrast with human PGC, which do not express Sox2 (16, 17, 15).

While the way towards quiescence involves the loss of proliferation and pluripotency markers, the data obtained here and in previous study (11) suggest that the transition from gonocytes to spermatogonia seems to depend on the upregulation of these factors. We have previously observed that the proliferation marker Ki67 and the pluripotent marker OCT4 are absent, at the protein level, from gonocytes from 19dpc to 5dpp. At around 8dpp these markers are back, suggesting that proliferation resumption and the reacquisition of pluripotent markers are linked (11). Conversely, here we detected SOX2 in 3dpp and 5dpp gonocytes, suggesting that this factor may have a function at this phase of rat male germ cell development. Interestingly, SOX2 was detected in the cytoplasm of these post-natal gonocytes, contrasting with the expected nuclear localization as a transcription factor. The cytoplasmic distribution of SOX2 in gonocytes showed a granulated pattern that is very similar to the processing bodies (P-bodies). Thus, we decided to look at the expression of the P-body marker GW182 in these cells. The P-bodies are subcellular structures where mRNAs and miRNAs are processed (26, 27, 28). The P-bodies contain proteins of the mRNA decay machinery and proteins involved in miRNA processing (29, 30, 31), indicating that the P-bodies participate of miRNA-mediated gene silencing. The protein GW182 interacts with Argonaute proteins and is fundamental for this process (32, 33). Here we observed that the labelling pattern of SOX2 and GW182 was very similar, suggesting that SOX2 is located to P-bodies in quiescent gonocytes and early spermatogonia. To confirm this hypothesis we performed the double-labelling of SOX2 and GW182, which is a key component of P-bodies and is related to gene silencing through miRNA pathway (24, 34, 32, 35, 36). SOX2 and GW182 co-localized in restricted portions of gonocytes cytoplasm. This leads to the hypothesis that SOX2 could play a role in mRNA processing at this phase of male germ cell development. Despite of the incomplete colocalization of SOX2 and GW182, we do not exclude a possible SOX2 function in RNA-processing since it could be part of distinct RNA-processing body in which GW182 is not present. As a positive evidence for this hypothesis, recent studies have shown that SOX2 interacts with RNA-processing proteins such as LIN28 (24) and RBM14 (37). Lin28, which is also associated with miRNA processing (38), localizes to P-bodies in undifferentiated embryonal carcinoma cells (39). Here we observed a partial co-localization of SOX2 and LIN28 in post-natal gonocyte cytoplasm, suggesting an interaction between these proteins at this phase male germ cell development. A partial interaction of SOX2 with LIN28 has been also observed in embryonic stem cells (24). On the other hand, most of LIN28 and GW182 protein detected in the gonocyte cytoplasm co-localized, suggesting that they are part of the same P-body class.

The nuclear detection of SOX2 protein as well as of Sox2 mRNA at 8dpp coincides with the return of Oct4 expression (11). Thus, our hypothesis is that SOX2 might be transported from the cytoplasm to the nucleus and that this coincides with the upregulation of OCT4. This could be an indicative of the emergence of the first spermatogonia in the rat testis, reinforcing our previous data suggesting that the first SSC emerge around 8dpp (11). Although *Nanog* has been detected only at mRNA level in the 8dpp testes, we consider that it represents another important indicative of spermatogonia emergence. Indeed, the frequency of spermatogonia showing nuclear SOX2 increases from 8dpp on so that, in the pre-pubertal testis, it was detected exclusively in the nucleus of rare undifferentiated spermatogonia. This data contrasts with humans in which the expression of SOX2 has not been detected in pre-pubertal testis (15). In mice, *Sox2* expression has been detected at mRNA level (19) but not at protein level (40) in adult testis.

In this study we described the dynamics of Sox2 expression in male germ cells from PGC stage to pre-pubertal spermatogonia stage. We conclude that by the end of mitotic arrest SOX2 protein is upregulated and locates to gonocytes cytoplasm to subsequently be transported to the nucleus, what happens concomitantly with *Nanog* and Oct4 re-expression. We also raise the hypothesis that SOX2 may have a function in RNA processing. Taking all these data together, we propose that SOX2 plays important and distinct roles in rat germ cell development and can be a good indicator of the reacquisition of the pluripotency potential by these cells.

## 6. ACKNOWLEDGEMENTS

The authors declare that there is no conflict of interest involved in this work. The authors thank Fundacao de Amparo a Pesquisa do Estado de Sao Paulo (FAPESP, Proc. No. 2012/08951-1) and CAPES for financial support. The authors also thank Isabelle Hernandez Cantão and Ana Flavia Popi for critical reading of the manuscript.

## 7. REFERENCES

1. C. Huckins: C: The spermatogonial stem cell population in adult rats. I. Their morphology, proliferation and maturation. Anat Rec 169(3):533–57 (1971)

2. R. Ollinger, A.J. Childs, H.M. Burgess, R.M. Speed, P.R. Lundegaard, N. Reynolds, N.K. Gray, H.J. Cooke, I.R. Adams: Deletion of the pluripotency-associated Tex19.1. gene causes activation of endogenous retroviruses and defective spermatogenesis in mice. PLoS Genet 4(9):e1000199 (2008)

3. K. Gassei, K.E. Orwig: SALL4 expression in gonocytes and spermatogonial clones of postnatal mouse testes. PLoS One 8(1):e53976 (2013)

4. N. Aeckerle, K. Eildermann, C. Drummer, J. Ehmcke, S. Schweyer, A. Lerchl, M. Bergmann, S. Kliesch, J. Gromoll, S. Schlatt, R. Behr: The pluripotency factor LIN28 in monkey and human testes: a marker for spermatogonial stem cells? Mol Hum Reprod 18(10):477–88 (2012)

5. K. Zheng, X. Wu, K.H. Kaestner, P.J. Wang: The pluripotency factor LIN28 marks undifferentiated spermatogonia in mouse. BMC Dev Biol 9:38 (2009)

6. M. Pesce, X. Wang, D.J. Wolgemuth, H. Schöler: Differential expression of the Oct-4 transcription factor during mouse germ cell differentiation. Mech Dev 71(1-2):89–98 (1998)

7. M. Kanatsu-Shinohara, K. Inoue, J. Lee, M. Yoshimoto, N. Ogonuki, H. Miki, S. Baba, T. Kato, Y. Kazuki, S. Toyokuni, M. Toyoshima, O. Niwa, M. Oshimura, T. Heike, T. Nakahata, F. Ishino, A. Ogura, T. Shinohara: Generation of pluripotent stem cells from neonatal mouse testis. Cell 119:1001–12 (2004)

8. K. Guan, K. Nayernia, L.S. Maier, S. Wagner, R. Dressel, J.H. Lee, J. Nolte, F. Wolf, M. Li, W. Engel, G. Hasenfuss: Pluripotency of spermatogonial stem cells from adult mouse testis. Nature 440:1199–203 (2006)

9. F. Izadyar, F. Pau, J. Marh, N. Slepko, T. Wang, R. Gonzalez, T. Ramos, K. Howerton, C. Sayre, F. Silva: Generation of multipotent cell lines from a distinct population of male germ line stem cells. Reproduction 135:771–784 (2008)

10. M. Culty: Gonocytes, from the fifties to the present: is there a reason to change the name? Biol Reprod 89(2):46 (2013)

11. C. Zogbi, R.B. Tesser, G. Encinas, S.M. Miraglia, T. Stumpp: Gonocyte development in rats: proliferation, distribution and death revisited. Histochem Cell Biol 138:305–22 (2012)

12. P.S. Western, J.A. van den Bergen, D.C. Miles, A. H. Sinclair: Male fetal germ cell differentiation involves complex repression of the regulatory network controlling pluripotency. FASEB J 24(8):3026–35 (2010)

13. R.T. Mitchell, G. Cowan, K.D. Morris, R.A. Anderson, H.M. Fraser, K.J. Mckenzie, W.H. Wallace, C.J. Kelnar, P.T. Saunders, R.M. Sharpe: Germ cell differentiation in the marmoset (Callithrix jacchus) during fetal and neonatal life closely parallels that in the human. Hum Reprod 23(12):2755–65 (2008)

14. C.E. Hoei-Hansen, K. Almstrup, J.E. Nielsen, S.B. Sonne, N. Graem, N.E. Skakkebaek, H. Leffers, E. Rajpert-De Meyts: Stem cell pluripotency factor NANOG is expressed in human fetal gonocytes, testicular carcinoma in situ and germ cell tumours. Histopathology 47(1):48–56 (2005)

15. S.B. Sonne, R.M. Perrett, J.E. Nielsen, M.A. Baxter, D.M. Kristensen, H. Leffers, N.A. Hanley, E. Rajpert-De-Meyts: Analysis of SOX2 expression in developing human testis and germ cell neoplasia. Int J Dev Biol 54(4):755–60 (2010)

16. J. de Jong, H. Stoop, A.J. Gillis, R.J. van Gurp, G.J. van de Geijn, M.D. Boer, R. Hersmus, P.T. Saunders, R.A. Anderson, J.W. Oosterhuis, L.H. Looijenga Differential expression of SOX17 and SOX2 in germ cells and stem cells has biological and clinical implications. J Pathol 215(1):21–30 (2008)

17. R.M. Perrett, L. Turnpenny, J.J. Eckert, M. O’Shea, S.B. Sonne, I.T. Cameron, D.I. Wilson, E. Rajpert-De Meyts, N.A. Hanley: The early human germ cell lineage does not express SOX2 during in vivo development or upon in vitro culture. Biol Reprod 78(5):852–8 (2008)

18. A.A Ketkar, K.V.R. Reddy: Expression Pattern of OCT-4 and PLZF Transcription Factors during the Early Events of Spermatogenesis in Mice. J Cell Sci Ther 3:120 (2012)

19. M. Imamura, K. Miura, K. Iwabuchi, T. Ichisaka, M. Nakagawa, J. Lee, M. Kanatsu-Shinohara, T. Shinohara, S. Yamanaka: Transcriptional repression and DNA hypermethylation of a small set of ES cell marker genes in male germline stem cells. BMC Dev Biol 6:34 (2006)

20. S. Yamaguchi, H. Kimura, M. Tada, N. Nakatsuji, T. Tada: Nanog expression in mouse germ cell development. Gene Expr Patterns 5(5):639–46 (2005)

21. D. Bhartiya, S. Kasiviswanathan, S.K. Unni, P. Pethe, J.V. Dhabalia, S. Patwardhan, H.B. Tongaonkar: Newer Insights Into Premeiotic Development of Germ Cells in Adult Human Testis Using Oct-4 as a Stem Cell Marker. J Histochem Cytochem 58(12):1093–1106 (2010)

22. G. Encinas, C. Zogbi, T. Stumpp: Detection of four germ cell markers in rats during testis morphogenesis: differences and similarities with mice. Cells Tissues Organs 195:443–55 (2012)

23. H.M. Beaumont, A.M. Mandl: A quantitative study of primordial germ cells in the male rat. J Embryol Exp Morphol 11:715–40 (1963)

24. J.L. Cox, S.K. Mallanna, X. Luo, A. Rizzino: Sox2 uses multiple domains to associate with proteins present in Sox2-protein complexes. PLoS One 5(11):e15486 (2010)

25. P.S. Western, D.C. Miles, J.A. van den Bergen, M. Burton, A.H. Sinclair: Dynamic regulation of mitotic arrest in fetal male germ cells. Stem Cells 26:339–47 (2008)

26. P. Anderson, N. Kedersha: RNA granules. J Cell Biol 172:803–808 (2006)

27. A. Eulalio, I. Behm-Ansmant, E. Izaurralde: P bodies: at the crossroads of posttranscriptional pathways Nat Rev Mol Cell Biol 8:9–22 (2007)

28. R. Parker, U. Sheth: P bodies and the control of mRNA translation and degradation. Mol Cell 18. 25:635–46 (2007)

29. A. Jakymiw, K.M. Pauley, S. Li, K. Ikeda, S. Lian, T. Eystathioy, M. Satoh, M.J. Fritzler, E.K. Chan: The role of GW/P-bodies in RNA processing and silencing. J Cell Sci 120(Pt 8):1317–23. A. Jakymiw, K.M. Pauley, S. Li, K. Ikeda, S. Lian, T. Eystathioy, M. Satoh, M.J. Fritzler, E.K. Chan Erratum in: J Cell Sci. 2007;120(Pt 9):1702 (2007)

30. S.P. Chan, F.J. Slack: microRNA-mediated silencing inside P-bodies. RNA Biol 3(3):97–100 (2006)

31. J. Liu, M.A. Valencia-Sanchez, G.J. Hannon, R. Parker: MicroRNA-dependent localization of targeted mRNAs to mammalian P-bodies. Nat Cell Biol 7(7):719–23 (2005)

32. A. Eulalio, E. Huntzinger, E. Izaurralde: GW182 interaction with Argonaute is essential for miRNA-mediated translational repression and mRNA decay. Nat Struct Mol Biol 15(4):346–53 (2008)

33. J. Pfaff, J. Hennig, F. Herzog, R. Aebersold, M. Sattler, D. Niessing, G. Meister: Structural features of Argonaute-GW182 protein interactions. Proc Natl Acad Sci U S A 110(40):E3770–9 (2013)

34. L. Ding, M. Han: GW182 family proteins are crucial for microRNA-mediated gene silencing. Trends Cell Biol 17(8):411–6 (2007)

35. A. Eulalio, F. Tritschler, E. Izaurralde:. The GW182 protein family in animal cells: new insights into domains required for miRNA-mediated gene silencing. RNA 15(8):1433–42 (2009)

36. J.E. Braun, E. Huntzinger, E. Izaurralde: The role of GW182 proteins in miRNA-mediated gene silencing. Adv Exp Med Biol 768:147–63 (2013)

37. X Fang, J.G. Yoon, L. Li, Y.S. Tsai, S. Zheng, L. Hood, D.R. Goodlett, G. Foltz, B. Lin: Landscape of the SOX2 protein-protein interactome. Proteomics 11(5):921–34 (2011)

38. S.R. Viswanathan, G.Q. Daley, R.I. Gregory: Selective blockade of microRNA processing by Lin28. Science 320(5872):97–100 (2008)

39. E. Balzer, E.G. Moss: Localization of the developmental timing regulator Lin28 to mRNP complexes, P-bodies and stress granules. RNA Biol 4(1):16–25 (2007)

40. K. Arnold, A. Sarkar, M.A. Yram, J.M. Polo, R. Bronson, S. Sengupta, M. Seandel, N. Geijsen, K. Hochedlinger: Sox2(+) adult stem and progenitor cells are important for tissue regeneration and survival of mice. Cell Stem Cell 9(4):317–29 (2011)

